# ModkitOpt: Accurate RNA modification site calling from nanopore sequencing

**DOI:** 10.64898/2025.12.19.695383

**Authors:** Alexandra Sneddon, Stefan Prodic, Eduardo Eyras

**Affiliations:** EMBL Australia Partner Laboratory Network at the Australian National University, Canberra ACT 2601, Australia; Centre for Computational Biomedical Sciences, The John Curtin School of Medical Research, Australian National University, Canberra ACT 2601, Australia; The Shine-Dalgarno Centre for RNA Innovation, The John Curtin School of Medical Research, Australian National University, Canberra ACT 2601, Australia; The Australian Research Council Centre of Excellence for the Mathematical Analysis of Cellular Systems, Australia

## Abstract

**Summary:** ModkitOpt is a pipeline for the systematic optimisation of modkit filtering thresholds and stoichiometry cutoffs using orthogonally validated modification sites. It substantially improves site-calling accuracy across modification types, even when the nanopore sample and validation sites are from different biological contexts. We report optimal site-calling parameters for m6A and pseU and provide practical recommendations for applying ModkitOpt.

**Availability and Implementation:** ModkitOpt is freely available under the MIT license at https://github.com/comprna/modkitopt.

**Supplementary Information:** Supplementary data is available at the journal’s web site.

## Introduction

Nanopore direct RNA sequencing enables single-molecule detection of RNA modifications at single-nucleotide resolution^1^. Yet the identification of modification sites across the transcriptome relies on ad hoc processing of per-read predictions^2^, risking inaccurate site calling and reduced reproducibility.

A typical workflow estimates modification stoichiometry (fraction of molecules modified) at each transcriptomic site by aggregating per-read predictions from Oxford Nanopore Technologies’ (ONT) dorado^3^ using ONT’s modkit^4^ *pileup*. Sites are then classified as modified or unmodified using a stoichiometry cutoff. Consequently, site-calling accuracy depends on two critical steps: (1) modkit’s stoichiometry estimation, and (2) appropriate cutoff selection.

Despite its widespread use, modkit’s parameters have not been systematically evaluated for their impact on site-calling accuracy. Modkit provides two thresholds to filter low-confidence canonical (*--filter-threshold*) and modified (*--mod-threshold*) base predictions. By default, these are estimated heuristically without calibration against site-calling accuracy, potentially increasing false discoveries or excluding true modification sites^5^. This may particularly affect rare modifications such as pseudouridine (pseU), for which thresholds can be biased towards the more abundant canonical predictions. Although users can override these defaults, threshold choice relies on subjective judgement or ad hoc experimentation.

Stoichiometry cutoff selection introduces further uncertainty. Despite its central role in defining modification landscapes, no standardised or validated method exists for selecting a cutoff that optimises false positive and false negative rates.

Here, we systematically evaluate the impact of modkit parameters and stoichiometry cutoff on site-calling accuracy across multiple RNA modification types. We introduce ModkitOpt, a framework that optimises modkit parameters and stoichiometry cutoff for a given dataset and modification type using orthogonally validated sites, providing an efficient and reproducible workflow for nanopore RNA modification site calling. We also report optimal site-calling parameters for m6A and pseU and provide practical recommendations for applying ModkitOpt.

### Site-calling parameter sensitivity

To evaluate the impact of modkit thresholding on site-calling performance, we compared nanopore-derived modification sites against orthogonally validated GLORI-derived m6A^6^ and BACS-derived pseU^7^ sites from HEK293T and HeLa cells, respectively. For both modifications, modkit’s default thresholding strategy was substantially less accurate than most tested threshold combinations (**Figure S1a-b**). Performance was driven primarily by the modified base threshold (*--mod_threshold*), with more stringent filtering improving precision at the expense of recall. Optimal thresholds differed between modifications, with pseU requiring more stringent filtering than m6A.

Since modkit filters low-confidence dorado predictions before stoichiometry estimation, we investigated whether site-calling threshold sensitivity was related to dorado prediction confidence. For m6A, dorado assigned substantially higher confidence predictions to GAC motifs, the most abundant m6A motif (**Figure S2**), than to other sequence contexts (**Figure S3**). Restricting analysis to GAC sites largely eliminated threshold sensitivity, reducing the performance difference between default and optimal thresholds to 2.0% (**Figure S4**), compared with 53% across all sequence contexts (**Figure S1a**). This suggests that modkit threshold optimisation is most critical when dorado prediction confidence is heterogeneous.

Finally, using the optimal modkit thresholds, we assessed the impact of stoichiometry cutoff selection. For both modifications, increasing the cutoff increased precision at the expense of recall (**Figures S1c-d**), highlighting the need for principled cutoff optimisation.

### ModkitOpt implementation

To address both sources of uncertainty, we developed ModkitOpt (**Figure 1a**), a computational framework that jointly optimises modkit filtering thresholds and corresponding stoichiometry cutoff using validated modification sites. Given a modBAM file and validated sites, ModkitOpt quantifies precision, recall and F1 score across 36,000 parameter combinations and identifies the optimal configuration. While orthogonally validated sites are supplied for mammalian m6A (GLORI-derived; HEK293T^6^) and pseU (BACS-derived; HeLa^7^), users can also provide validated sites for any other modification type supported by ONT, enabling ModkitOpt’s broad application.

**Figure 1.**
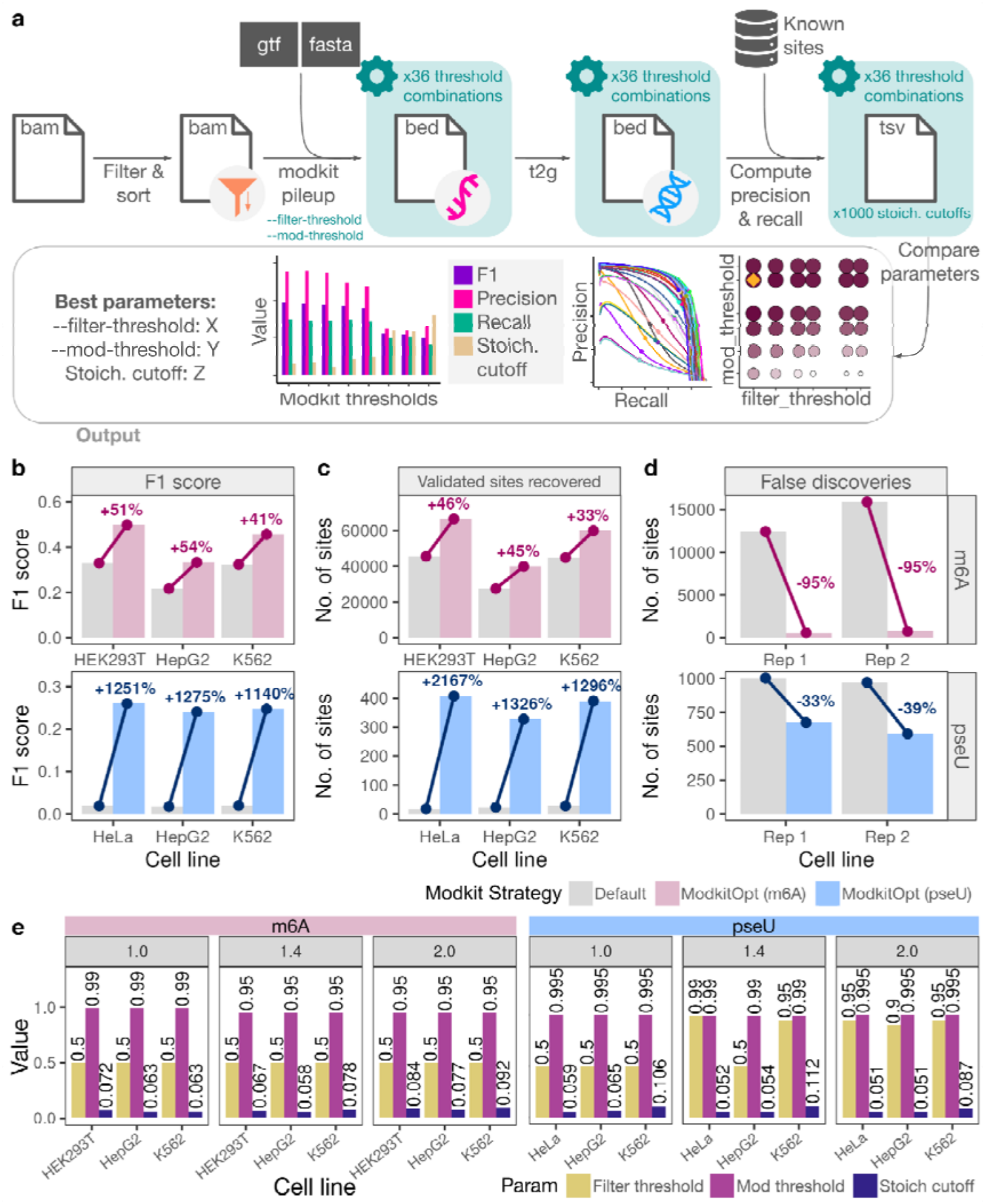
**a**, Overview of ModkitOpt. Per-read modification calls are processed with modkit *pileup* across a grid of *--filter-threshold* and *--mod-threshold* values to estimate site-level stoichiometry. Predicted sites are benchmarked against validated sites across 1,000 stoichiometry cutoffs, before the optimal modkit thresholds and stoichiometry cutoff, selected by F1 score, are reported alongside plots depicting site-calling performance across the parameter space (Figures S6-8). **b-d**, Performance of ModkitOpt (pink, N6-methyladenosine (m6A); blue, pseudouridine (pseU)) versus default modkit with an optimised stoichiometry cutoff (grey), measured by F1 score (**b**), validated called sites (**c**) and false-positive sites recovered from non-modified IVT controls (**d**). The number of evaluable sites (coverage ≥ 20 reads) for each IVT dataset and filtering strategy is provided in **Table S3**. Validated sites: GLORI m6A sites (HEK293T) and BACS pseU sites (HeLa). **e**, Optimal *--filter-threshold* (yellow), *--mod-threshold* (purple) and stoichiometry cutoff (indigo) identified by ModkitOpt for m6A and pseU across dorado versions (panel titles).

#### Pipeline

ModkitOpt requires a transcriptome-aligned modBAM, reference transcriptome (FASTA), gene annotation (GTF or GFF3), and modification type (e.g., m6A, pseU, m5C or inosine). After filtering to retain primary alignments, the modBAM is sorted and indexed using samtools^8^ (v1.21). Modkit^4^ (v0.6.0) *pileup* is then run across all combinations of *--filter-threshold* and *--mod-threshold* values (0.5, 0.75, 0.9, 0.95, 0.99, 0.995), producing transcriptome-coordinate bedMethyl files containing site-level stoichiometry estimates. Users can optionally specify additional threshold values to evaluate.

Predicted sites are converted to genomic coordinates for comparison with validated sites, requiring exact single-nucleotide coordinate matches. All validated sites are included in performance evaluation to avoid introducing sample-specific preprocessing (**Figure S5**). Where multiple transcriptomic positions map to the same genomic site, stoichiometry estimates are aggregated using a valid-coverage-weighted average. ModkitOpt evaluates 1,000 stoichiometry cutoffs uniformly spaced between 0 and 0.999, retaining sites with stoichiometry exceeding each cutoff.

Precision (fraction of predicted sites in the validated set), recall (fraction of validated sites predicted) and F1 score 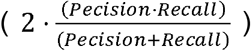 are calculated for every parameter combination. For each combination, the stoichiometry cutoff that maximises the F1 score is identified, and the overall parameter combination achieving the highest F1 is reported.

#### Outputs

ModkitOpt reports the optimal *--filter-threshold, --mod-threshold* and stoichiometry cutoff, together with the corresponding bedMethyl file, which the user can filter using the optimal cutoff. Additional outputs include plots and a TSV file with the precision, recall and F1 scores across the parameter space (**Figures S6-S8**).

#### Resource allocation and execution time

ModkitOpt is distributed as a Nextflow workflow for reproducible execution across computing environments, with automatic resource allocation. The default resource settings are: 8 CPUs and 8GB RAM for samtools filter and samtools sort; 8 CPUs and 30GB RAM for modkit *pileup*; 4 CPUs and 8GB RAM for samtools index; 1 CPU, 30GB RAM and 5GB job filesystem disk space for converting transcriptomic to genomic coordinates; and 1 CPU and 8GB RAM for all other tasks. Using these settings (**Supplementary Methods**), execution on an HPC system (PBSPro Scheduler) took 22 minutes for a 7GB modBAM file and 33 minutes for a 21GB modBAM file (**Table S2**).

### ModkitOpt site-calling performance

ModkitOpt consistently outperformed the default site-calling strategy of applying modkit with its heuristically estimated filtering parameters. For a conservative comparison, when assessing performance for the default strategy, we used the default modkit filtering parameters but independently optimised the stoichiometry cutoff for each nanopore dataset, selecting the cutoff that maximised F1 score. Precision, recall and F1 score were evaluated across stoichiometry cutoffs to identify the optimal cutoff for the default strategy for each dataset (**Figure S9**). Using orthogonally validated sites, ModkitOpt substantially improved site-calling performance across all nanopore datasets per modification, in terms of F1 score (**Figure 1b**), precision and recall (**Figure S10**). These improvements were maintained even when validated sites and the nanopore sample were from different biological contexts, suggesting that incomplete overlap between validation and sample-specific modification landscapes is sufficient for effective parameter optimisation. Of note, predicted sites in the nanopore sample that are not in the validated sites are not necessarily false positives; thus, the precision and F1 scores reported here should be interpreted as reflecting the agreement with the available orthogonally validated sites rather than absolute accuracy.

We next investigated the robustness of ModkitOpt parameter optimisation to limited overlap between nanopore samples and validated sites. Remarkably, even when optimisation was restricted to just 5% of the validated sites shared with the nanopore sample, ModkitOpt achieved median relative F1 score improvements of 44% (range: 23-61%) for m6A and 434% (range: 198-1866%) for pseU (**Figure S11**). These improvements reflected gains in both precision and recall relative to default modkit parameters, although the larger relative improvement for pseU is influenced by its lower F1 score under default parameters. Greater improvements were observed when 10% or more of the shared sites were used for optimisation. Importantly, the optimal *--mod-threshold*, the most influential modkit parameter (**Figure S1**), remained largely consistent for m6A (0.95-0.995) and pseU (0.99-0.995) across overlap sizes down to 5% (**Figure S12**). Given the substantial variation in modification site overlap between tissues (**Tables S4-S7**), these results demonstrate that ModkitOpt can achieve robust parameter optimisation even when only a small proportion of validated sites are shared with the nanopore sample, supporting its applicability across diverse biological contexts.

To evaluate the generalisability of ModkitOpt’s parameter optimisation, we froze the optimal parameters identified by ModkitOpt and reassessed site-calling performance for the same nanopore datasets but using independent reference datasets. Performance gains were recapitulated across reference datasets (**Figures S13 and S14**). Interestingly, for each modification type, the optimal modkit threshold combinations were identical across all datasets (**Table S1**). Although this may suggest that the optimal thresholds could reflect the modification’s frequency and stoichiometry, these properties can also change between conditions. Importantly, ModkitOpt enables parameter optimisation independently per dataset, without additional assumptions.

Optimisation of site-calling parameters with ModkitOpt substantially altered the inferred RNA modification landscapes. ModkitOpt consistently recovered validated sites that were missed by the default strategy (**Figures 1c and S10b**), including up to 22-fold more validated pseU sites. Importantly, these gains were accompanied by a decrease infalse predictions, estimated as modification calls in fully non-modified in vitro transcribed (IVT) RNA controls^9^ (**Figures 1d and S15**). While the validated sites do not necessarily reflect the full modification landscape of a nanopore sample, preventing assessment of absolute site calling accuracy, these findings demonstrate that site-calling parameter selection can profoundly affect reported RNA modification landscapes and suggest that previously reported landscapes may benefit from re-evaluation using optimised parameters.

Finally, we identified the optimised site-calling parameters for different dorado versions (**Figures 1e and S16**), finding that for m6A the optimal *--filter-threshold* is consistently 0.5 for all versions, while the optimal *--mod-threshold* is 0.99 for tested versions up to v1.0, and 0.95 for newer versions. For pseU, the optimal *--mod-threshold* is 0.995 for all tested versions except v1.4, for which 0.99 is optimal. The best *--filter-threshold* is 0.5 for tested versions up to v1.0, with 0.9-0.99 most commonly best for newer versions. The optimal stoichiometry cutoff varied between 0.06-0.09 for m6A and 0.05-0.1 for pseU. Considering our previous analysis that indicated ModkitOpt performance is robust down to a 5% overlap of sites between the nanopore sample and validated sites, these optimal parameters are expected to be broadly applicable. However, when the overlap between the nanopore sample and validated dataset represents less than 5% of the sites in either dataset, users are recommended to run ModkitOpt using a more representative orthogonal dataset, if available.

## Conclusions

Accurate identification of RNA modification sites is essential for translating nanopore direct RNA sequencing into reliable biological insight. Here, we show that RNA site-calling accuracy is highly sensitive to both modkit filtering parameters and stoichiometry cutoff selection, and that modkit’s default heuristic parameters can substantially alter inferred modification landscapes. ModkitOpt addresses these challenges by systematically optimising both steps using validated modification sites, consistently improving site-calling performance across modification types, datasets and biological contexts. We recommend using the dorado-version-specific optimised parameters identified here when at least 5% of sites in both the nanopore sample and orthogonal validation dataset are shared; below this, ModkitOpt should be applied using a more representative orthogonal validation dataset, where available. As nanopore-based modification detection continues to mature, standardised and validated post-processing will be critical to ensure reproducible and comparable modification landscapes. ModkitOpt achieves this through an efficient and intuitive workflow, allowing standardised evaluation of both existing and future RNA modification studies. ModkitOpt is available at https://github.com/comprna/modkitopt.

## Acknowledgements

We acknowledge indirect support by the Australian Government’s National Collaborative Research Infrastructure Strategy (NCRIS) through access to computational resources provided by the National Computational Infrastructure (NCI). We are grateful to the personnel from the Biomolecular Resource Facility at JCSMR (ANU) for their technical assistance and continued support regarding ONT sequencing. EE and AS are members of the Human RNome Consortium.

## Availability of data and materials

ModkitOpt is available at https://github.com/comprna/modkitopt (https://doi.org/10.5281/zenodo.24295546). The nanopore datasets for HeLa, HepG2 and K562 cells (RNA004) that were analysed during the current study are available from the European Nucleotide Archive (ENA) under accession PRJEB80229 (sample accessions SAMEA116055276 for HepG2, SAMEA116055277 for K562 and SAMEA116055274 for HeLa). The nanopore dataset for HEK293T cells (RNA004) is available from the SG-NEx project at registry.opendata.aws/sg-nex-data” (SGNex_Hek293T_directRNA_replicate5_run1). The nanopore datasets from unmodified IVT RNA from human hippocampus samples are available from ENA under accession PRJEB106788 (sample accessions SAMEA120997328 for Rep1 and SAMEA120997329 for Rep2). The GLORI-derived validated sites for m6A in HEK293T and HeLa cells were obtained from Supplementary Data 1 and Supplementary Data 5, respectively, of Liu et al. (Nature Biotechnology 2022), while the BACS-derived and PRAISE-derived validated sites for pseU were obtained from Supplementary Table 8 of Xu et al. (Nature Methods 2024), and from Supplementary Dataset 2 of Zhang et al. (Nature Chemical Biology 2023), respectively.

## Author contributions

AS, SP and EE conceptualised the study. AS led the software development and data analysis, with essential contributions from SP & EE. AS wrote the manuscript. SP and EE reviewed the manuscript.

## References

1. Garalde, D. R. et al. Highly parallel direct RN A sequencing on an array of nanopores. Nat. Methods 15, 201–206 (2018).

2. Cruciani, S. & Novoa, E. M. The new era of single-molecule RNA modification detection through nanopore base-calling models. Nature Reviews Molecular Cell Biology 2025 27:1 27, 10–18 (2025).

3. nanoporetech/dorado: Oxford Nanopore’s Basecaller. https://github.com/nanoporetech/dorado.

4. nanoporetech/modkit: A bioinformatics tool for working with modified bases. https://github.com/nanoporetech/modkit.

5. Zou, Y. et al. A comparative evaluation of computational models for RNA modification detection using nanopore sequencing with RNA004 chemistry. Brief. Bioinform. 26, (2025).

6. Liu, C. et al. Absolute quantification of single-base m6A methylation in the mammalian transcriptome using GLORI. Nat. Biotechnol. 41, 355–366 (2023).

7. Xu, H. et al. Absolute quantitative and base-resolution sequencing reveals comprehensive landscape of pseudouridine across the human transcriptome. Nat. Methods 21, 2024–2033 (2024).

8. Li, H. et al. The Sequence Alignment/Map format and SAMtools. Bioinformatics 25, 2078–2079 (2009).

9. Prodic, S. et al. SWARM resolves nanopore signal interference between RNA modification types and reveals splicing-shaped pseudouridylation. bioRxiv 2025.12.18.695332 (2026) doi:10.64898/2025.12.18.695332.

