## Supplementary Material for "ModkitOpt: Accurate RNA modification site calling from nanopore sequencing"

### Supplementary Methods

#### Default site-calling performance evaluation

Impact of modkit filtering thresholds

Site-calling performance was assessed by comparing nanopore-derived modification sites from HEK293T^1^ and HeLa^2^ cells with orthogonally validated sites generated using chemical conversion-based methods in the corresponding cell lines – GLORI (m6A, HEK293T)^3^ and BACS (pseU, HeLa)^4^. Nanopore pod5 files were processed using dorado^5^ (v0.8.3) (*dorado basecaller rna004_130bps_sup@v5.1.0 $fasta --modified-bases-models $model --reference $reference --mm2-opts “-k 14”^6^*) with m6A model rna004_130bps_sup@v5.1.0_inosine_m6A@v1, pseU model rna004_13bps_sup@v5.1.0_pseU@v1 and the human reference transcriptome GRCh38.p14 release 45^7^ to generate per-read modification predictions in modBAM files. The modBAM files were then filtered to retain primary alignments only, and sorted and indexed using samtools (v1.21)^8^. Modkit^9^ (v0.6.0) pileup was subsequently executed across all combinations of *--filter-threshold* and *--mod-threshold* values (0.5, 0.75, 0.9, 0.95, 0.99, 0.995), to produce transcriptome-coordinate bedMethyl files with site-level stoichiometry estimates. Sites were filtered for a minimum valid coverage (N_valid_cov_ output by modkit) of 20 reads, then converted to genomic coordinates using an R script (tidyverse^10^, GenomicFeatures^11^), with comparisons to validated sites requiring exact single-nucleotide coordinate matches. To account for multiple transcriptomic positions mapping to the same genomic site, stoichiometry values were aggregated using a valid-coverage-weighted average across transcripts. To isolate the effect of modkit filtering thresholds from downstream site-calling decisions, the stoichiometry cutoff was set to 0.1 for all threshold combinations.

Performance was measured using precision (fraction of the predicted sites that are in the validated set), recall (fraction of the validated sites that are predicted) and F1 score ($2\cdot\frac{\left( Pecision\cdot Recall \right)}{\left( Pecision+Recall \right)}$). Validated sites were restricted to those that had sufficient valid coverage (minimum 20 N_valid_cov_ output by modkit) in the respective nanopore dataset.

Impact of stoichiometry cutoff

Using stoichiometry estimates generated with the modkit threshold combination that achieved highest F1 score above for m6A in HEK293T cells and pseU in HeLa cells, site-calling performance was evaluated across 1,000 stoichiometry cutoffs uniformly spaced between 0 and 0.999. Sites exceeding each cutoff were classified as modified and compared against the validated site set. Precision, recall and F1 score were calculated as described above.

#### ModkitOpt performance evaluation

##### Site-calling performance

Site-calling performance of ModkitOpt was compared to the default site-calling strategy of applying modkit with its default heuristically-estimated filtering parameters. GLORI-derived^3^ m6A sites in HEK293T cells and BACS-derived^4^ pseU sites in HeLa cells were used as the reference dataset for ModkitOpt parameter optimisation. Nanopore datasets for HEK293T^1^, HepG2^2^ and K562^2^ cells were used to evaluate m6A site-calling performance, while nanopore datasets for HeLa^2^, HepG2^2^ and K562^2^ cells were used for pseU.

Nanopore pod5 files were parsed into modBAM files using dorado, as above, before applying ModkitOpt (setting --*fasta* and *--annotation* to the transcript sequences and comprehensive gene annotation, respectively, from the human reference transcriptome GRCh38.p14 release 45^7^). For a conservative comparison with ModkitOpt, we evaluated the default modkit site-calling strategy by applying the default modkit filtering parameters but independently optimising the stoichiometry cutoff for each nanopore dataset. Specifically, we evaluated 1,000 candidate stoichiometry cutoffs uniformly spaced between 0 and 0.999, and calculated precision, recall and F1 score at each cutoff, with the cutoff yielding the maximum F1 score selected for each dataset. The resulting performance curves and selected optimal cutoffs are shown in **Figure S9**. The performance of ModkitOpt and the default site-calling strategy was subsequently compared using F1 score and the number of validated sites recovered.

##### Effect of site overlap between nanopore and validated datasets

We evaluated the performance of ModkitOpt parameter optimisation when the number of shared sites between the nanopore dataset and validated sites dataset was progressively decreased. We first filtered a set of validated sites (GLORI-derived from HEK293T cells for m6A and BACS-derived from HeLa cells for pseU) to retain those with a valid coverage (N_valid_cov_ output by modkit) of at least 20 reads in a nanopore dataset (HEK293T cells for m6A and HeLa cells for pseU) (“covered validated sites”). 20% of these sites were randomly selected and reserved for performance evaluation. Parameter optimisation with ModkitOpt was performed using the remaining 80% of covered validated sites, as well as using randomly selected subsets of decreasing size (60%, 40%, 20%, 10% and 5%). The optimal parameters identified using each subset were then evaluated using the reserved test sites. For evaluation, validated sites that were identified in the nanopore dataset but were not part of the reserved test set were excluded from performance calculations to avoid falsely high reporting of false positives. This process of randomly splitting covered validated sites by a ratio of 80:20 for optimisation:test sets, and randomly selecting subsets of the optimisation set for parameter optimisation, was repeated 10 times with different random seeds to enable assessment of performance robustness to the choice of data split.

Importantly, since the validated sites provided by GLORI were in genomic coordinates and nanopore sites get converted by ModkitOpt to unique genomic coordinates such that multiple transcripts covering the same genomic coordinate are converted and aggregated into a single genomic site, there is no possibility for sites that are shared between transcript isoforms to appear in both optimisation and test sets. For conservative comparison of the default strategy to the ModkitOpt-optimised settings, the stoichiometry cutoff for the default strategy was optimised separately for each iteration (i.e., each combination of subset and seed), and the cutoff that maximised F1 score was selected. To compare the ModkitOpt-optimised settings with the default modkit implementation for each overlap size, we calculated the difference in performance between the two approaches for each random seed (i.e., paired comparisons using the same data split) and reported the median of these differences.

##### Evaluation on independent validated sites

The optimal parameters (--filter-threshold, --mod-threshold and stoichiometry cutoff) for each nanopore dataset were then frozen and site-calling performance was reassessed using a second reference dataset - GLORI-derived^3^ m6A sites in HeLa cells and PRAISE-derived^12^ pseU sites in HEK293T cells. Since PRAISE does not measure pseU with single-nucleotide resolution, to eliminate ambiguity in site comparisons, both PRAISE and the nanopore-derived sites were restricted to single-U sites.

##### False discovery analysis

Since it is not possible to determine whether unvalidated called sites in cellular-derived RNA are false discoveries, attributed to sample-specific modification landscapes, or limitations of the validated site measurement method, we instead used in vitro transcribed (IVT) RNA derived from human hippocampus samples^13^ as fully non-modified controls to evaluate false positive site calls. For each modkit parameter combination, the stoichiometry cutoff that was independently optimised on HEK293T cells for m6A and HeLa cells for pseU (as above in ”Site calling performance”) was applied. The default site-calling strategy was then compared against ModkitOpt using the optimised parameters identified for each modification type above. Given the IVTs are expected to be fully non-modified, all called sites were considered to be false discoveries.

##### Parameter optimisation per dorado version

We considered the 5 most recent dorado versions for which substantial changes to the RNA basecalling or modification models for m6A or pseU were introduced; 0.7 (new RNA basecalling models), 0.8 (new modification model), 1.0 (new basecalling and modification models), 1.4 (new basecalling models), 2.0 (new basecalling models and “high accuracy” modification models). Nanopore datasets for HEK293T^1^, HepG2^2^ and K562^2^ cells were used to evaluate m6A site-calling performance, while nanopore datasets for HeLa^2^, HepG2^2^ and K562^2^ cells were used for pseU. The following commands were used to run each dorado version:

- **v0.7.2:** dorado basecaller rna004_130bps_sup@v5.0.0 $pod5_dir --modified-bases <*m6A|pseU*>
- **v0.8.3:** dorado basecaller rna004_130bps_sup@v5.1.0 $pod5_dir --modified-bases-models rna004_130bps_sup@v5.1.0_inosine_m6A@v1,rna004_130bps_sup@v5.1.0_pseU@v1
- **v1.0.0:** dorado basecaller rna004_130bps_sup@v5.2.0 $pod5_dir --modified-bases-models rna004_130bps_sup@v5.2.0_inosine_m6A_2OmeA@v1,rna004_130bps_sup@v5.2.0_pseU_2OmeU@v1
- **v1.4.0:** dorado basecaller sup,pseU_2OmeU,inosine_m6A_2OmeA $pod5_dir
- **v2.0.1:** dorado basecaller sup,pseU_2OmeU,inosine_m6A_2OmeA $pod5_dir

Minimap2 was used to align each modBAM output to the reference transcriptome (GRCh38.p14 release 45) using options *-ax map-ont -uf -k 14 -t 32*. ModkitOpt was then run using modkit v0.6.0, the transcript sequences FASTA file and comprehensive gene annotation GFF3 file from the same reference transcriptome release used for minimap2, with additional ModkitOpt options --*mod_type m6A* and *–profile pbspro.*

##### Computational performance

The execution time of ModkitOpt was compared across 6 modBAM files ranging in size from 7.4GB to 21GB. Across all runs, ModkitOpt was executed on an HPC system (PBSPro scheduler) using default resource settings (8 CPUs and 8GB RAM for samtools filter and samtools sort; 8 CPUs and 30GB RAM for modkit pileup; 4 CPUs and 8GB RAM for samtools index; 1 CPU, 30GB RAM and 5GB job filesystem disk space for converting transcriptomic to genomic coordinates; and 1 CPU and 8GB RAM for all other tasks).

### Supplementary Figures

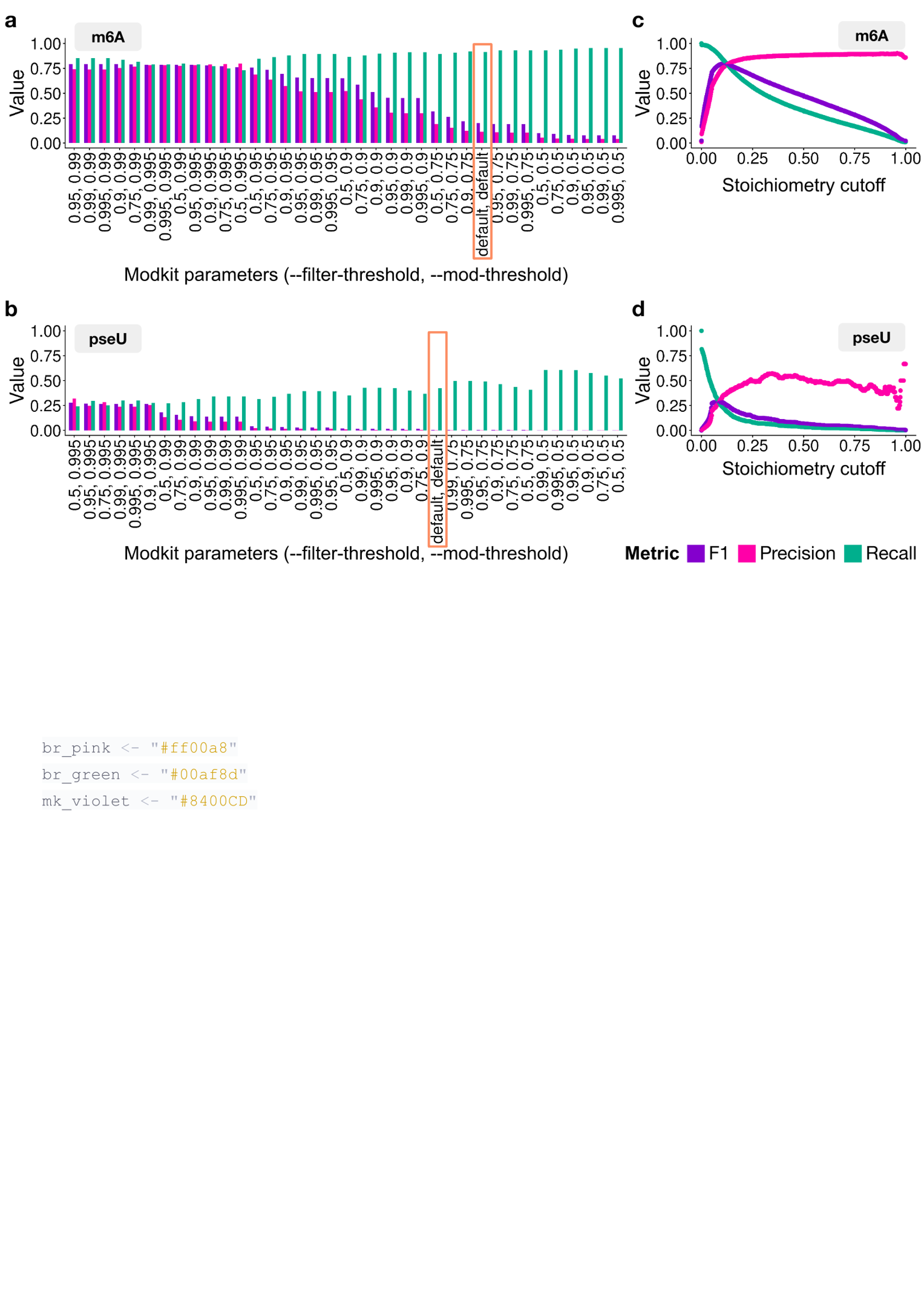

**Supplementary Figure S1:** **a-b,** Site-calling performance for N6-methyladenosine (m6A) in HEK293T cells (**a**) and pseudouridine (pseU) in HeLa cells (**b**) across 36 modkit threshold combinations, ordered by decreasing F1 score (purple), with precision (pink) and recall (green). The stoichiometry cutoff was fixed at 0.1. The modkit default is indicated (orange). **c-d,** Performance across stoichiometry cutoffs using the optimal threshold combination (highest F1 score) from (**a**) and (**b**), respectively**.** For this plot, recall was calculated only using validated sites (GLORI-derived sites from HEK293T cells for m6A or BACS-derived sites from HeLa cells for pseU) that had at least 20 reads valid coverage (N_valid_cov_ output by modkit) in the nanopore dataset.

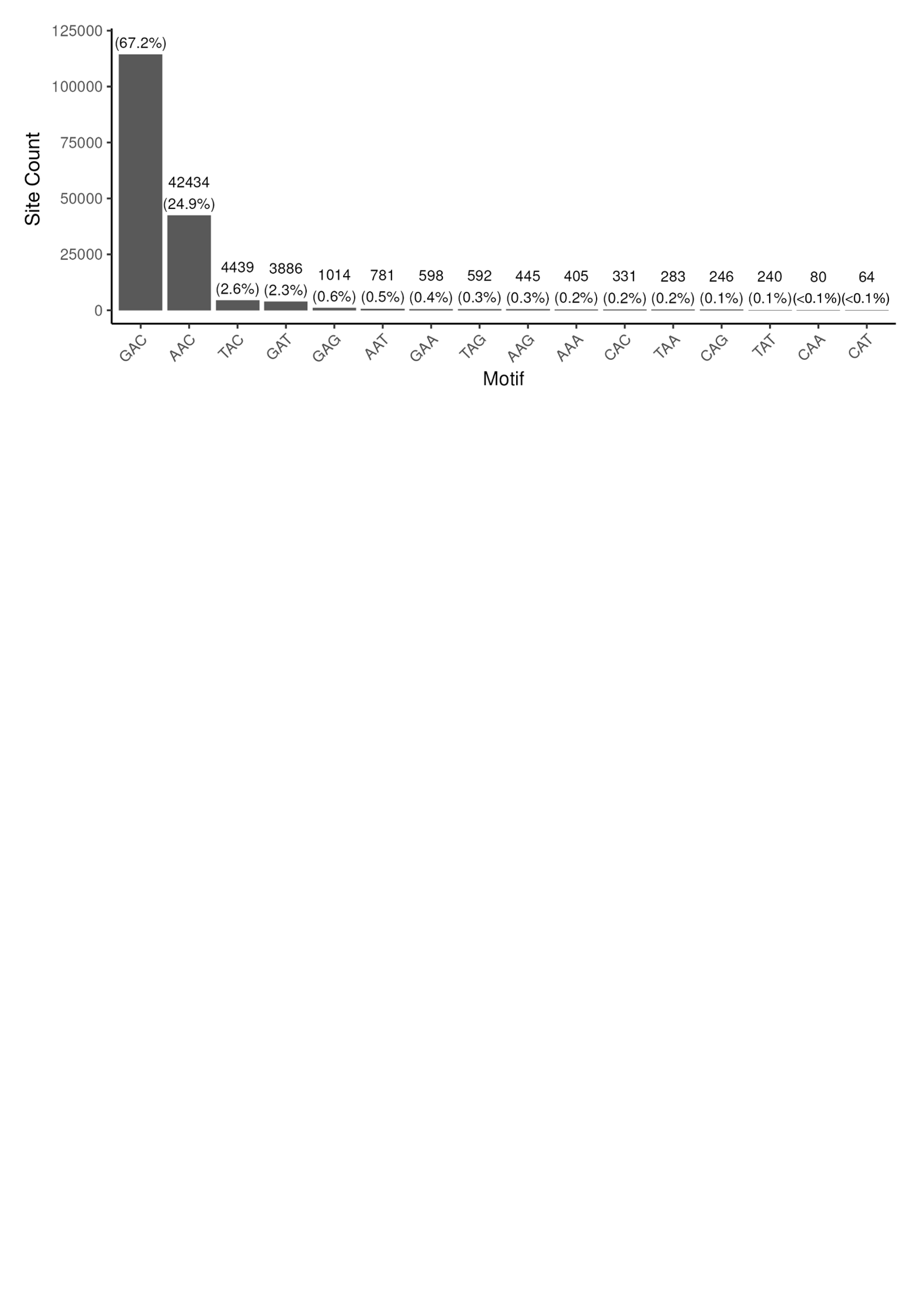

**Supplementary Figure S2:** Frequency of 3-mer sequence motifs centred on GLORI-defined N6-methyladenosine (m6A) sites. Motifs are ordered by abundance.

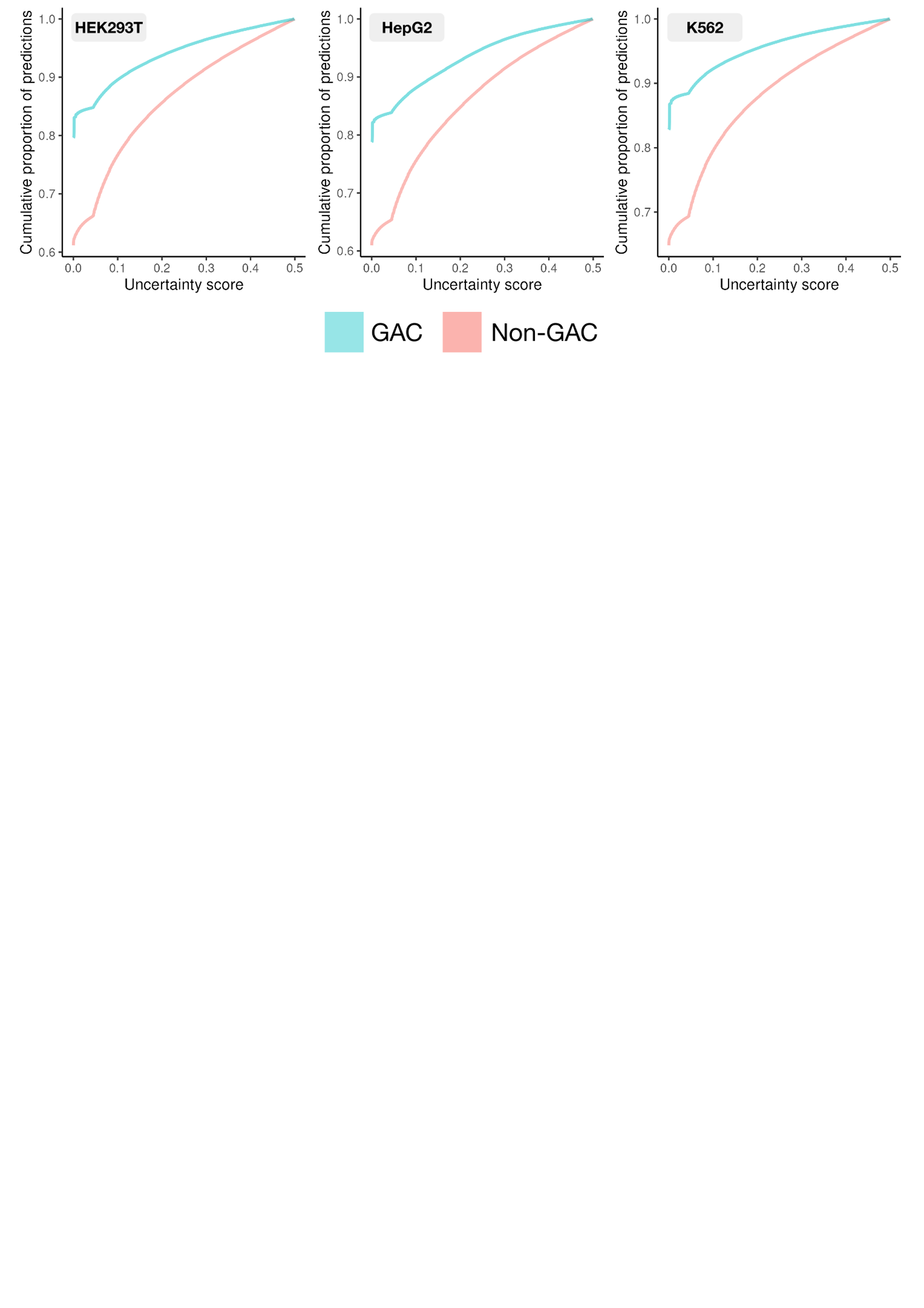

**Supplementary Figure S3:** Distribution of dorado prediction uncertainty in GAC (blue) and non-GAC (red) sequence contexts for nanopore datasets from HEK293T (left), HepG2 (centre) and K562 (right) cells. ModBAM files were generated using dorado (using m6A model rna004_130bps_sup@v5.1.0_inosine_m6A@v1, the human reference transcriptome GRCh38.p14 release 45, and options*-mm2-opts ”-k 14*”). For each dataset, 10% of m6A prediction probabilities at A sites were randomly sampled. Prediction uncertainty was defined as the minimum distance between each probability and 0 or 1, such that lower values correspond to higher-confidence predictions. Curves show the cumulative proportion of predictions as a function of increasing uncertainty.

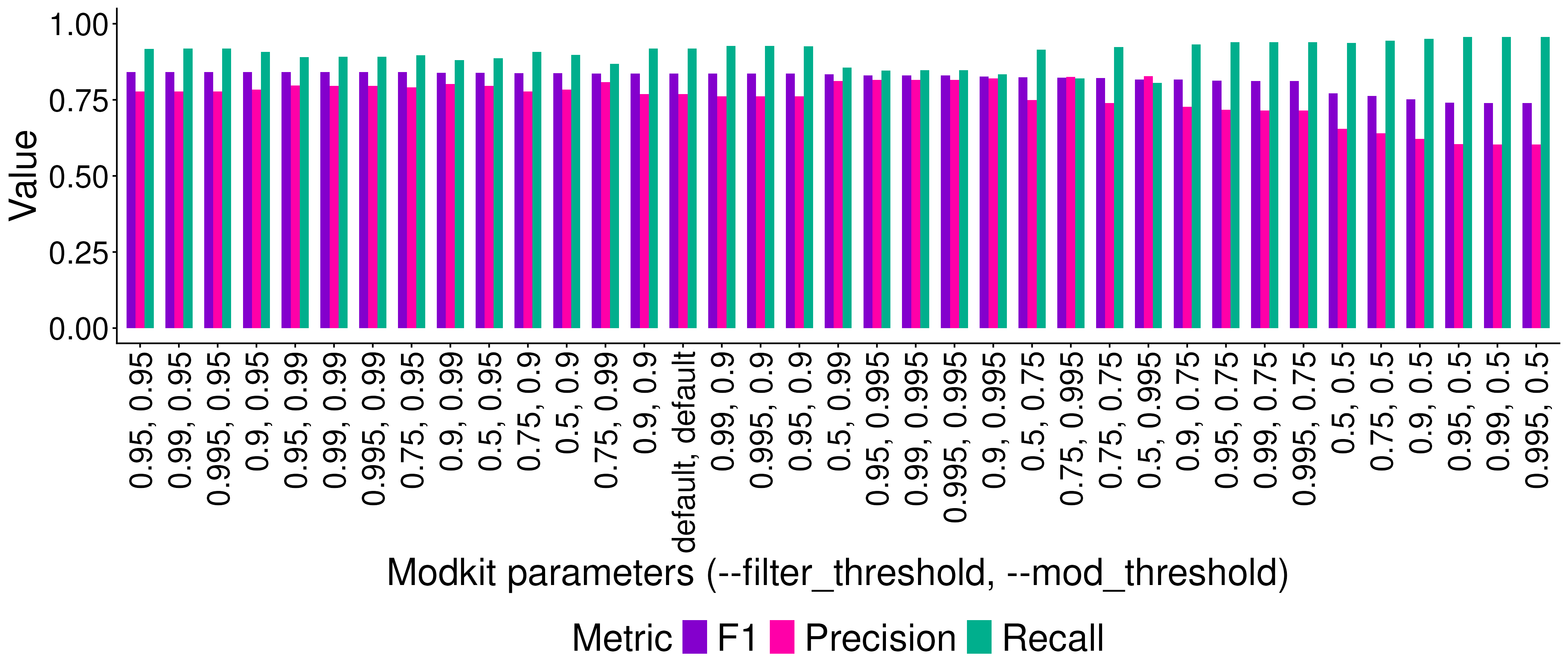

**Supplementary Figure S4:** Site-calling performance for GAC-centred N6-methyladenosine (m6A) sites in HEK293T cells across 36 modkit threshold combinations. Sites were restricted to the GAC motif using modkit’s *modbam adjust-mods --motif GAC*). Performance is shown as F1 score (purple)**,** precision (pink) and recall (green), with threshold combinations ordered by decreasing F1 score. For each combination, the stoichiometry cutoff was set to 0.1.

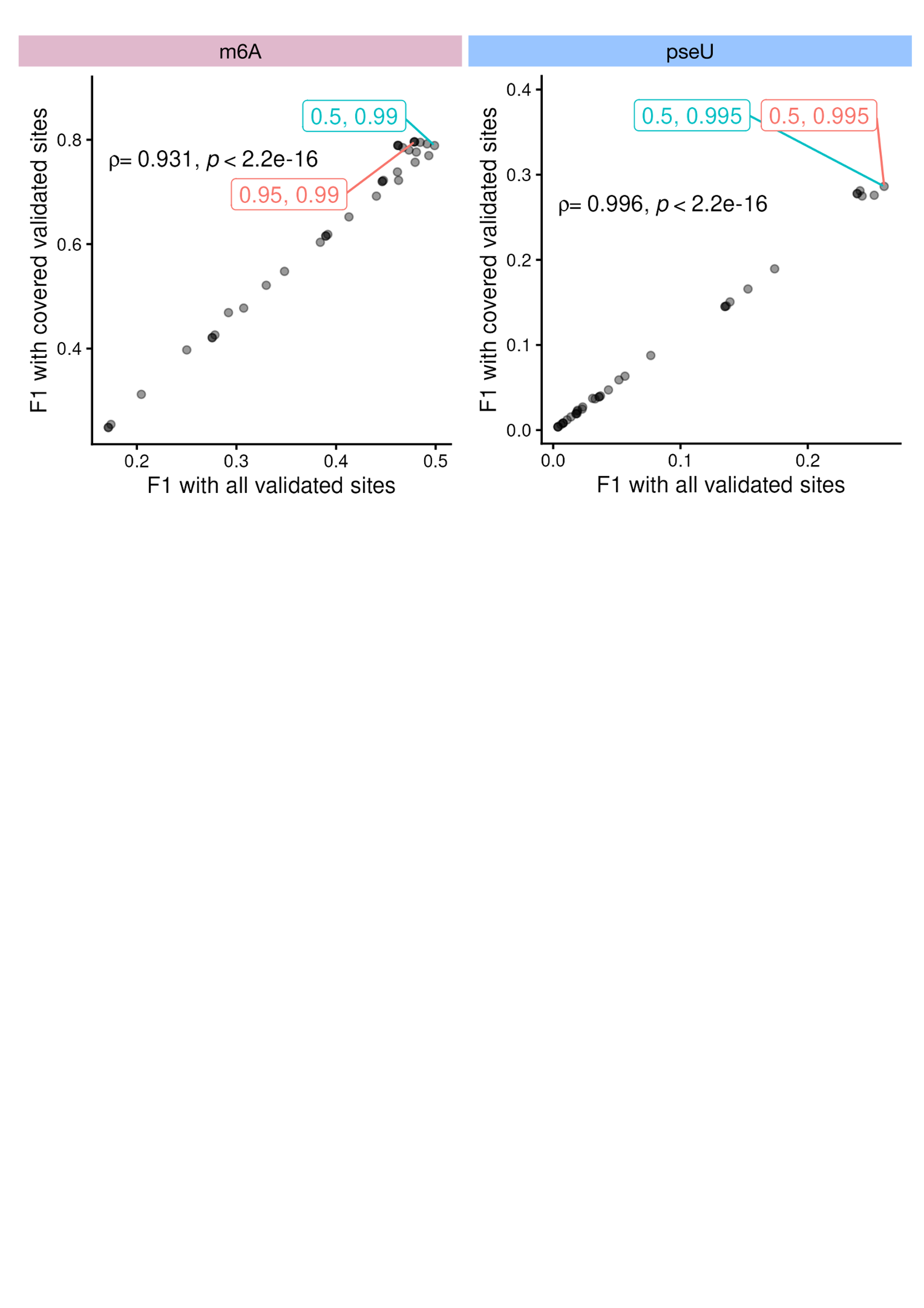

**Supplementary Figure S5:** Relationship between F1 scores calculated using as the denominator either all validated sites (x-axis) or only validated sites covered by the corresponding nanopore dataset (y-axis), quantified using Spearman’s rank-order correlation. The nanopore and validated sites datasets were from HEK293T cells for m6A (left), and HeLa cells (right) for pseU, with the validated sites derived from GLORI (HEK293T) and BACS (HeLa), respectively. Each point represents a modkit parameter combination, with the optimal parameter combinations for all validated sites and covered validated sites annotated in blue and red, respectively. Strong correlations between the two approaches indicate that the recall denominator has little effect on relative parameter rankings.

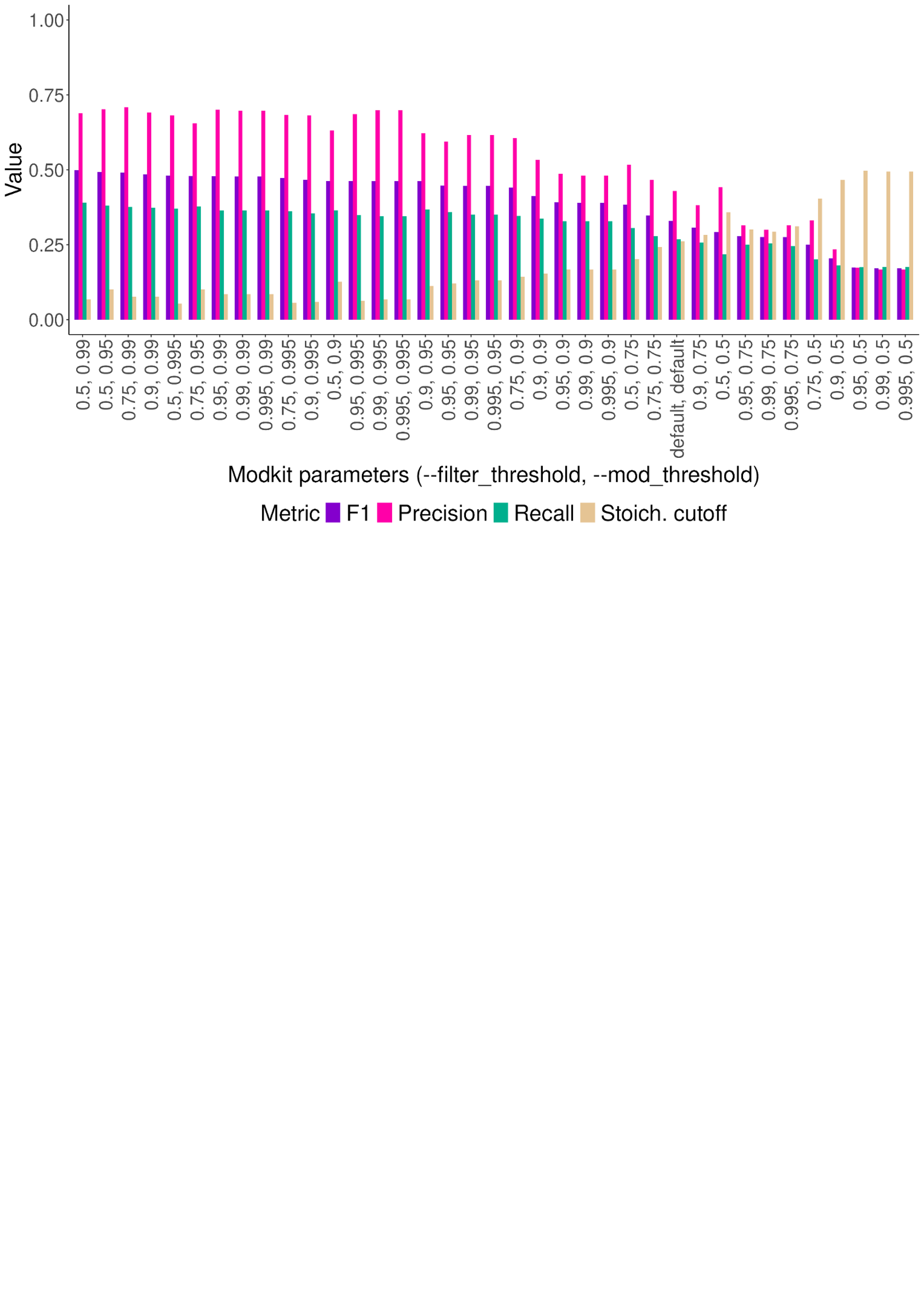

**Supplementary Figure S6**: Example plot output by ModkitOpt, showing the performance of modkit pileup across all combinations of --filter-threshold and --mod-threshold, sorted by decreasing F1 score, with modkit defaults indicated.

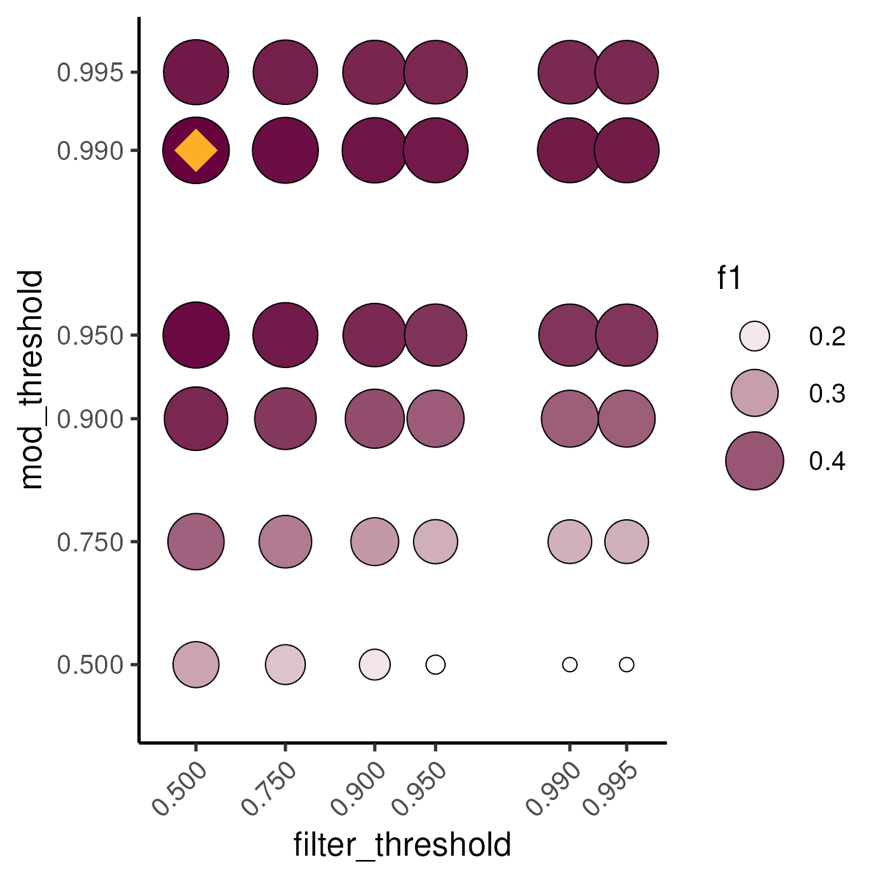

**Supplementary Figure S7**: Example plot output by ModkitOpt, showing the F1 score across all combinations of --filter-threshold (x-axis) and --mod-threshold (y-axis). For each threshold combination, the stoichiometry cutoff that maximised F1 score is shown. The optimal threshold combination is annotated (yellow diamond).

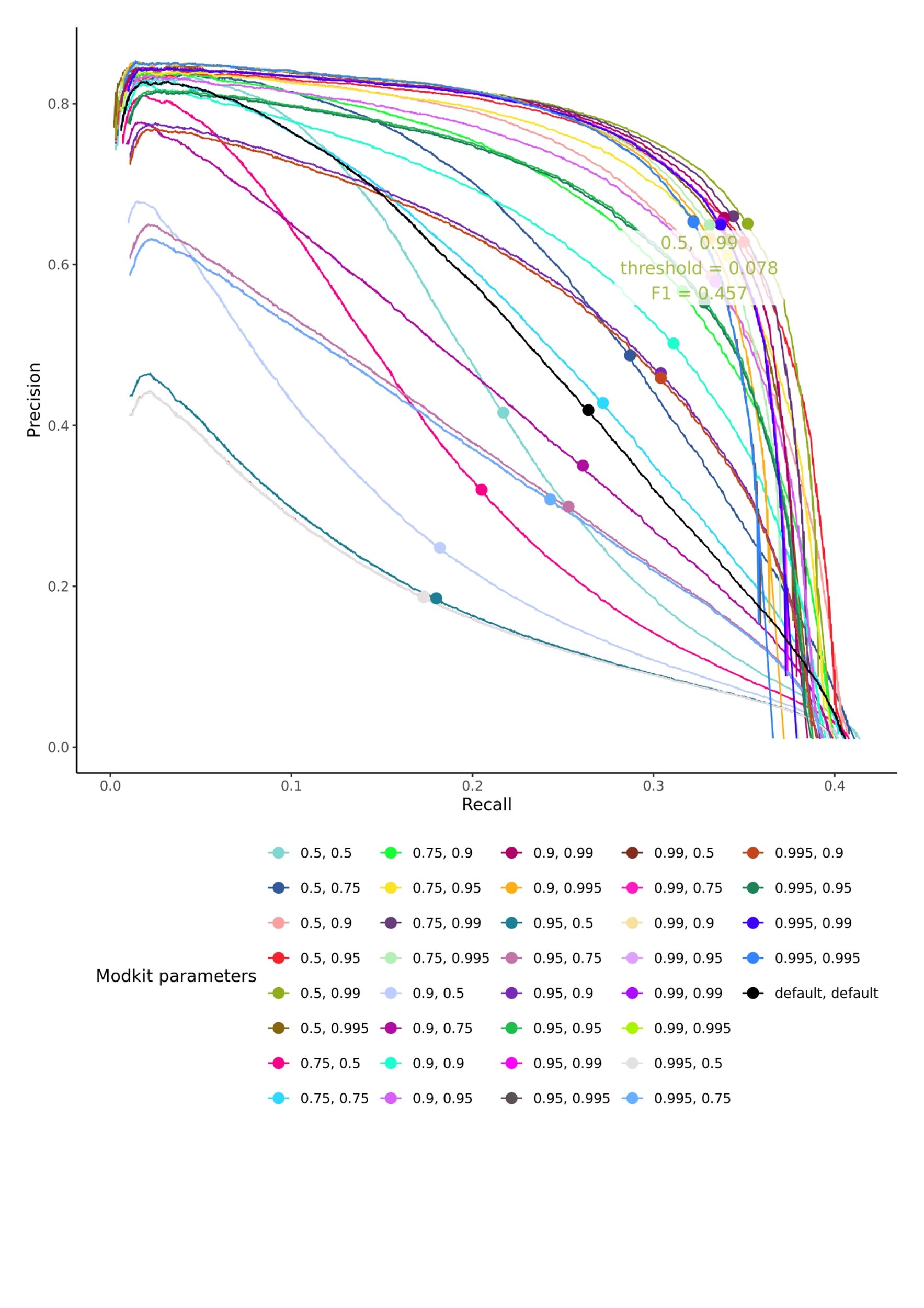

**Supplementary Figure S8**: Example plot output by ModkitOpt, showing the precision-recall curves across all combinations of --filter-threshold and --mod-threshold. For each threshold combination, the stoichiometry corresponding to the maximum F1 score is annotated with a circle.

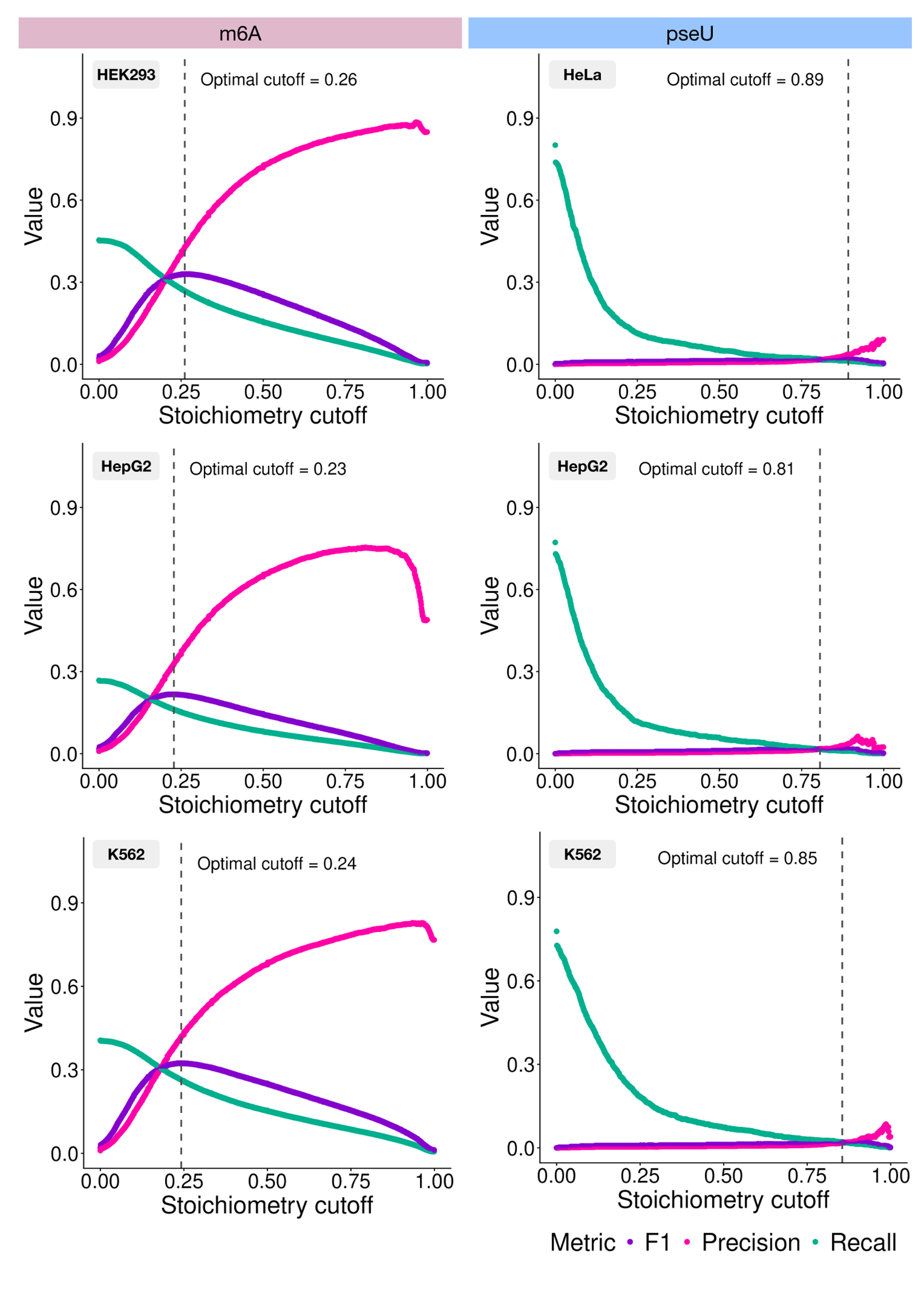

**Supplementary Figure S9:** Selection of optimal stoichiometry cutoffs for the default modkit site-calling strategy. F1 score (purple), precision (pink) and recall (green) are shown across stoichiometry cutoffs for m6A (left panel) and pseU (right panel). The default modkit filtering thresholds were used, with only the stoichiometry cutoff varied. The optimal cutoff, corresponding to the maximum F1 score, is indicated by a vertical dashed line and annotated for each cell line.

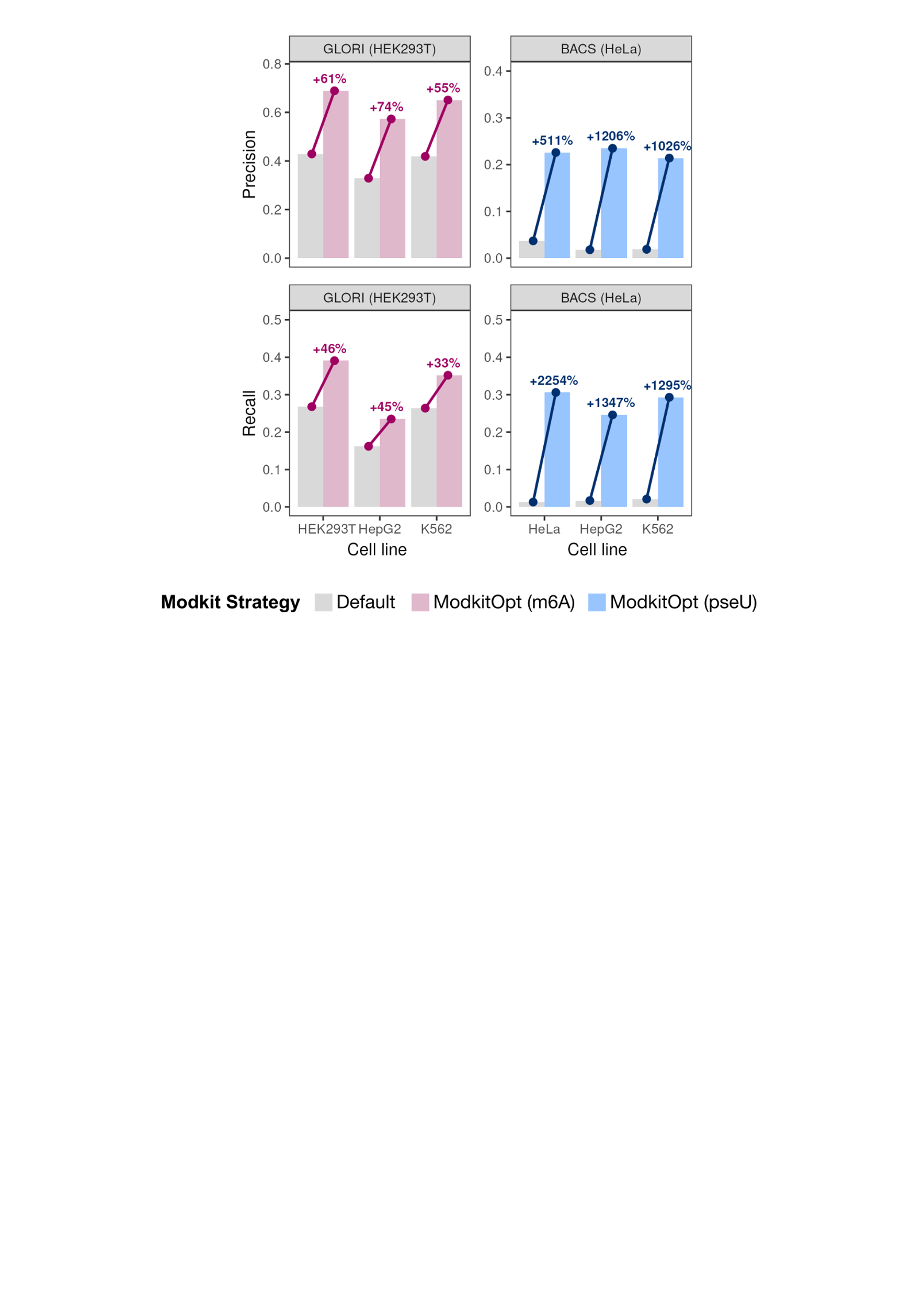

**Supplementary Figure S10:** Site-calling performance measured by precision (upper panels) and recall (lower panels) across nanopore datasets (x-axis) for N6-methyladenosine (m6A) (left) and pseudouridine (pseU) (right), comparing the default (grey) and ModkitOpt (coloured) site-calling strategies. Panel labels indicate the validated sites used for performance evaluation.

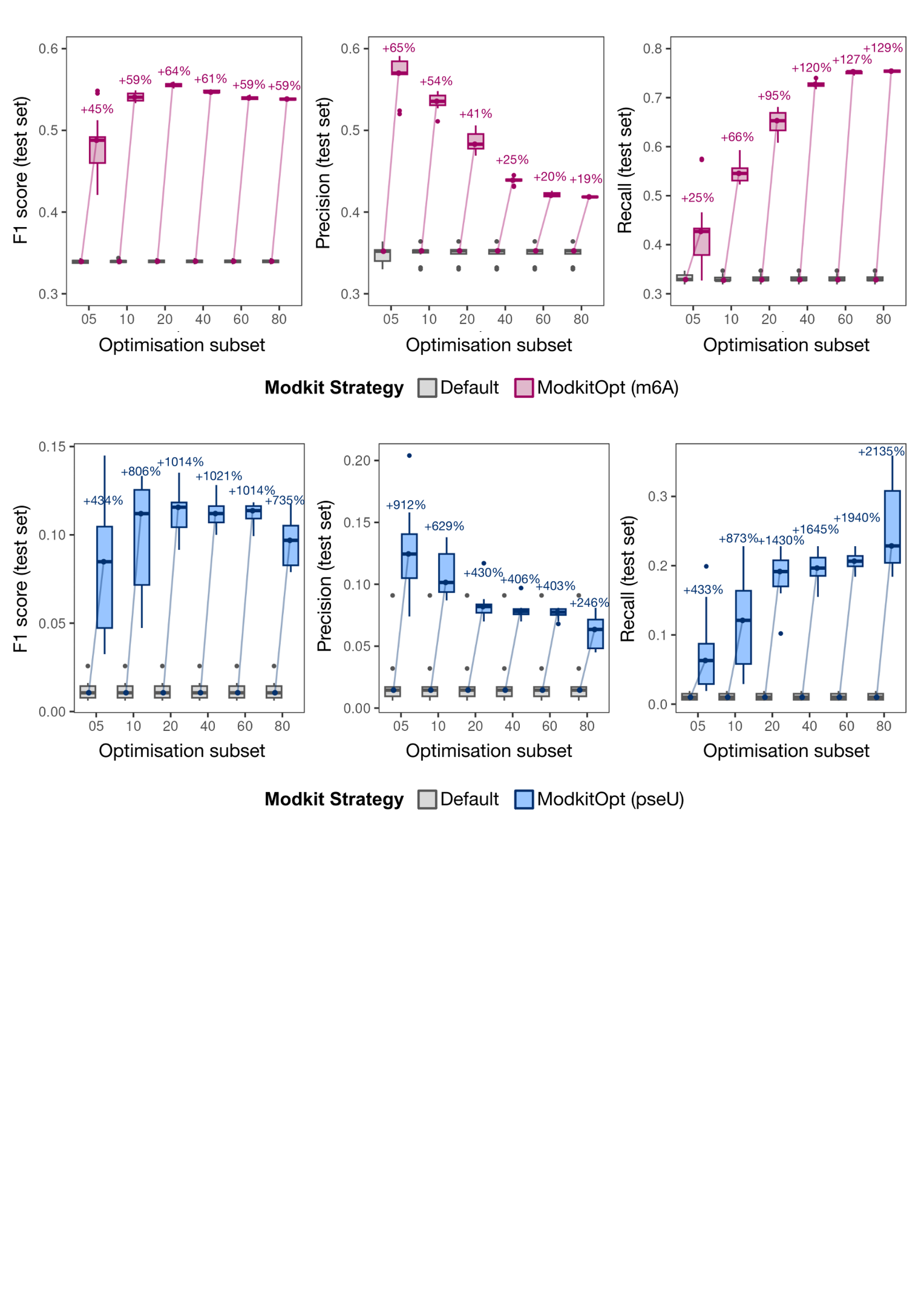

**Supplementary Figure S11**: Site-calling performance measured by F1 score (left), precision (middle) and recall (right) across increasing-sized subsets of optimisation sites (x-axis) for N6-methyladenosine (m6A) (upper) and pseudouridine (pseU) (lower), with each subset comprising a percentage of the total number of validated sites (GLORI-derived m6A in HEK293T cells and BACS-derived pseU in HeLa cells) that are covered by a nanopore dataset from HEK293T cells for m6A and HeLa cells for pseU. Performance was measured on the 20% of covered validated sites that were not used for parameter optimisation. Each optimisation:test split and random selection of site subsets was repeated 10 times with different random seeds. Performance is shown for the default modkit implementation (grey) and the ModkitOpt-optimised settings (magenta and blue), with bars connecting the median value between each pair of boxplots. Shown above each pair of boxplots is the median difference in performance between the default and ModkitOpt approaches for each random seed (i.e., the median of paired comparisons using the same data split).

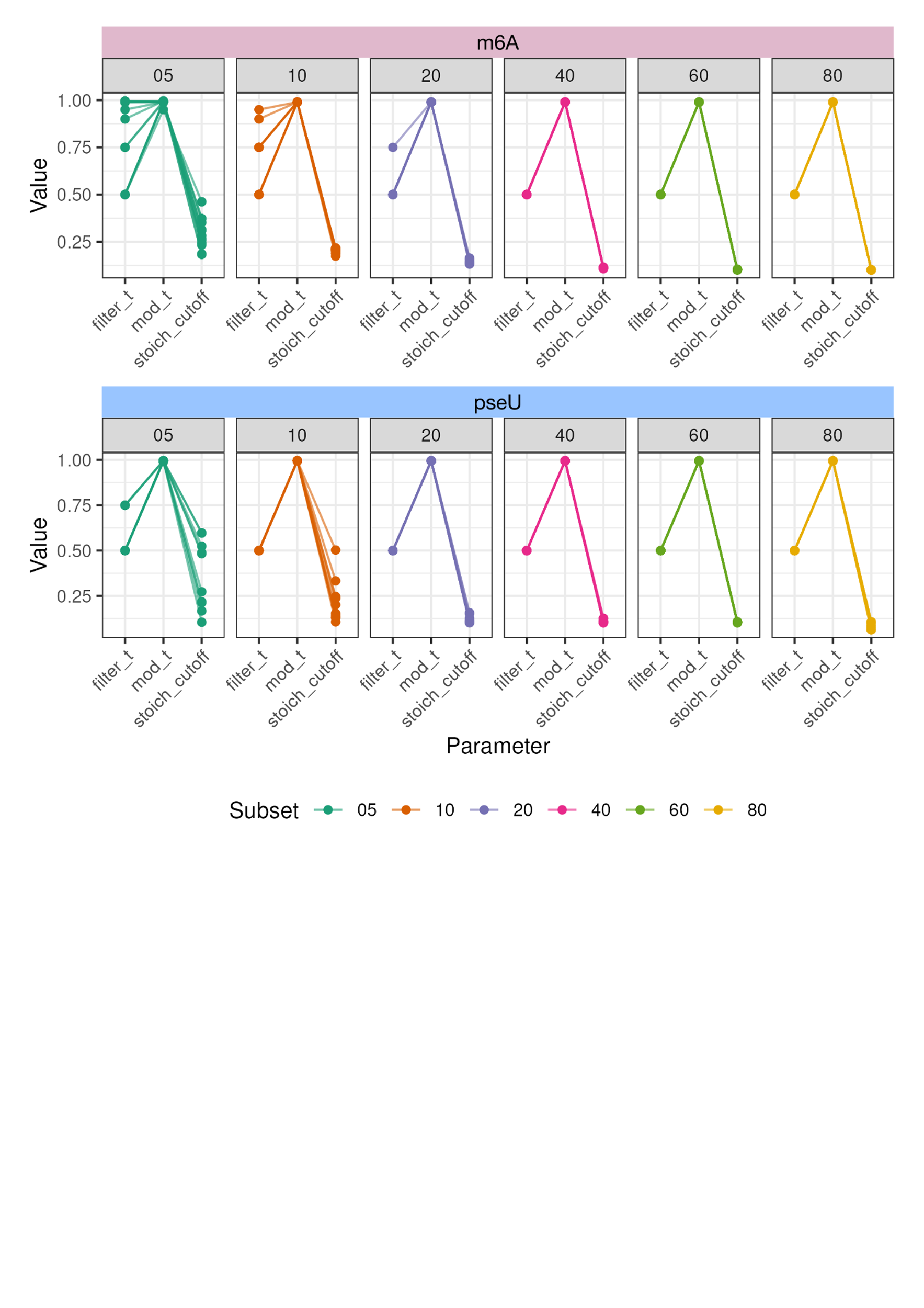

**Supplementary Figure S12**: Optimal parameters (filter_t = --filter-threshold, mod_t = --mod-threshold, stoich_cutoff = stoichiometry cutoff) identified by ModkitOpt for N6-methyladenosine (m6A) (upper) and pseudouridine (pseU) (lower) when parameter optimisation was performed using increasing-sized subsets of validated sites (panels labelled by subset size, as a percentage of the total number of validated sites that are covered by a nanopore dataset from HEK293T cells^1^ for m6A and HeLa cells for pseU). Each subset selection was repeated ten times with different random seeds. Validated sites were GLORI-derived m6A sites in HEK293T cells and BACS-derived pseU sites in HeLa cells. There are ten semi-transparent connected lines plotted per panel, each corresponding to a different seed used for subset selection.

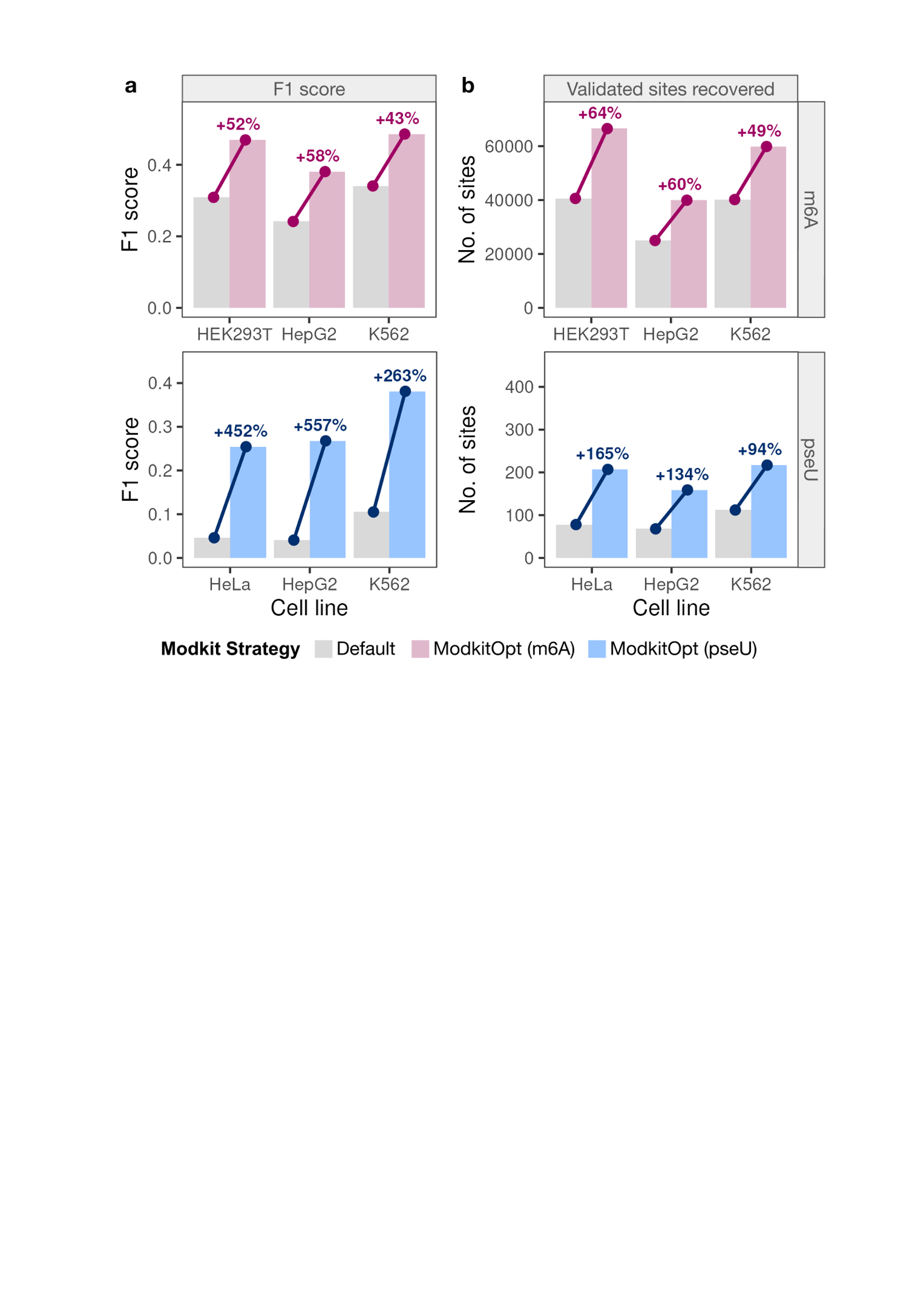

**Supplementary Figure S13:** Performance of ModkitOpt (pink, m6A; blue, pseU) versus default modkit with an optimised stoichiometry cutoff (grey), measured by F1 score (**a**) and validated called sites (**b**) across nanopore datasets (x-axis) for N6-methyladenosine (m6A) (upper panels) and pseudouridine (pseU) (lower panels). Validated sites were GLORI-identified in HeLa cells (m6A) and PRAISE-identified (single U sites only) in HEK293T cells (pseU).

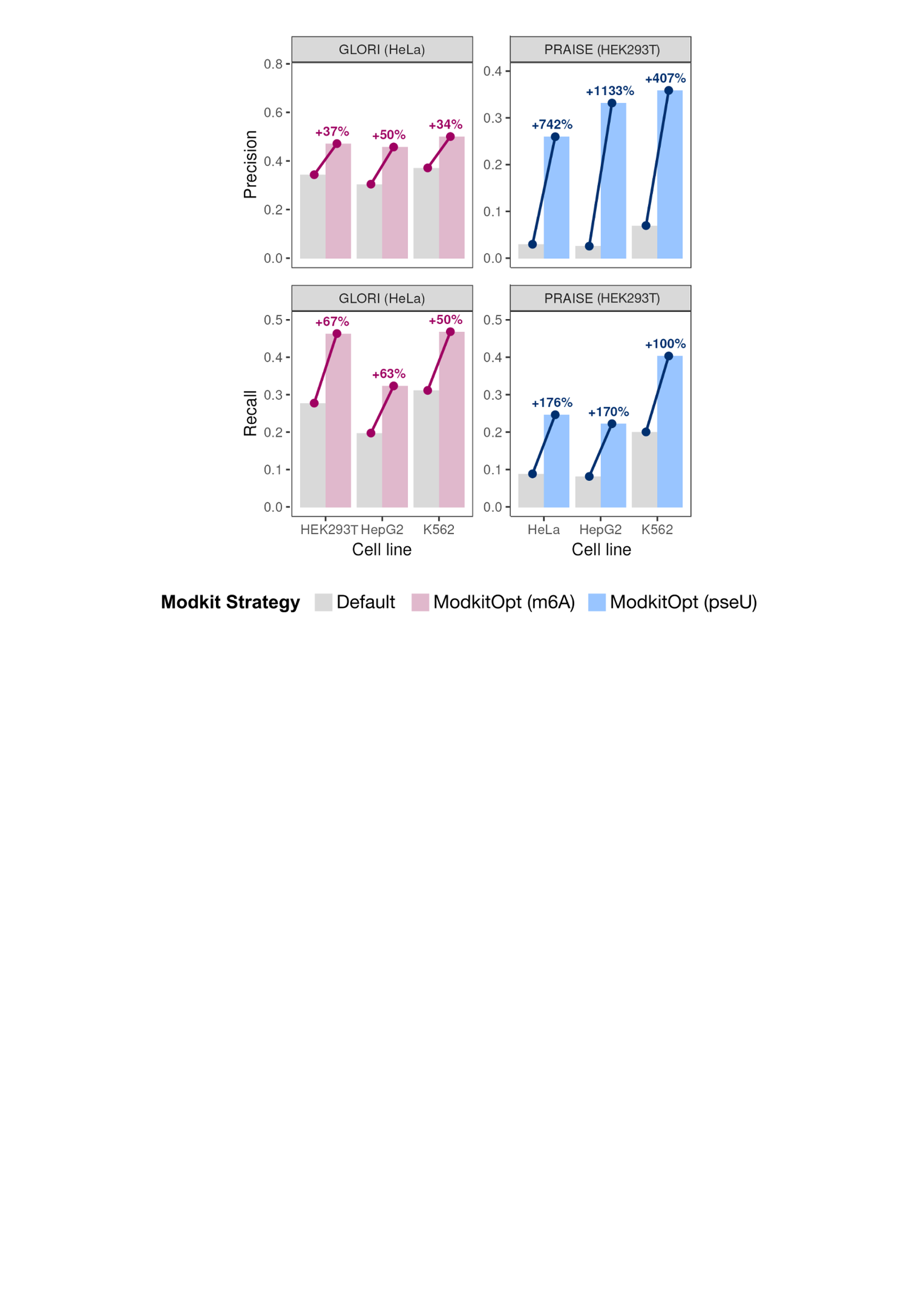

**Supplementary Figure S14:** Site-calling performance using alternative validated sites as a reference (indicated by panel labels) following parameter optimisation using default validated sites (GLORI-derived m6A sites in HEK293T cells and BACS-derived pseU sites in HeLa cells), measured by precision (upper panels) and recall (lower panels) across nanopore datasets (x-axis) for m6A (left) and pseU (right), comparing the default (grey) and ModkitOpt (coloured) site-calling strategies.

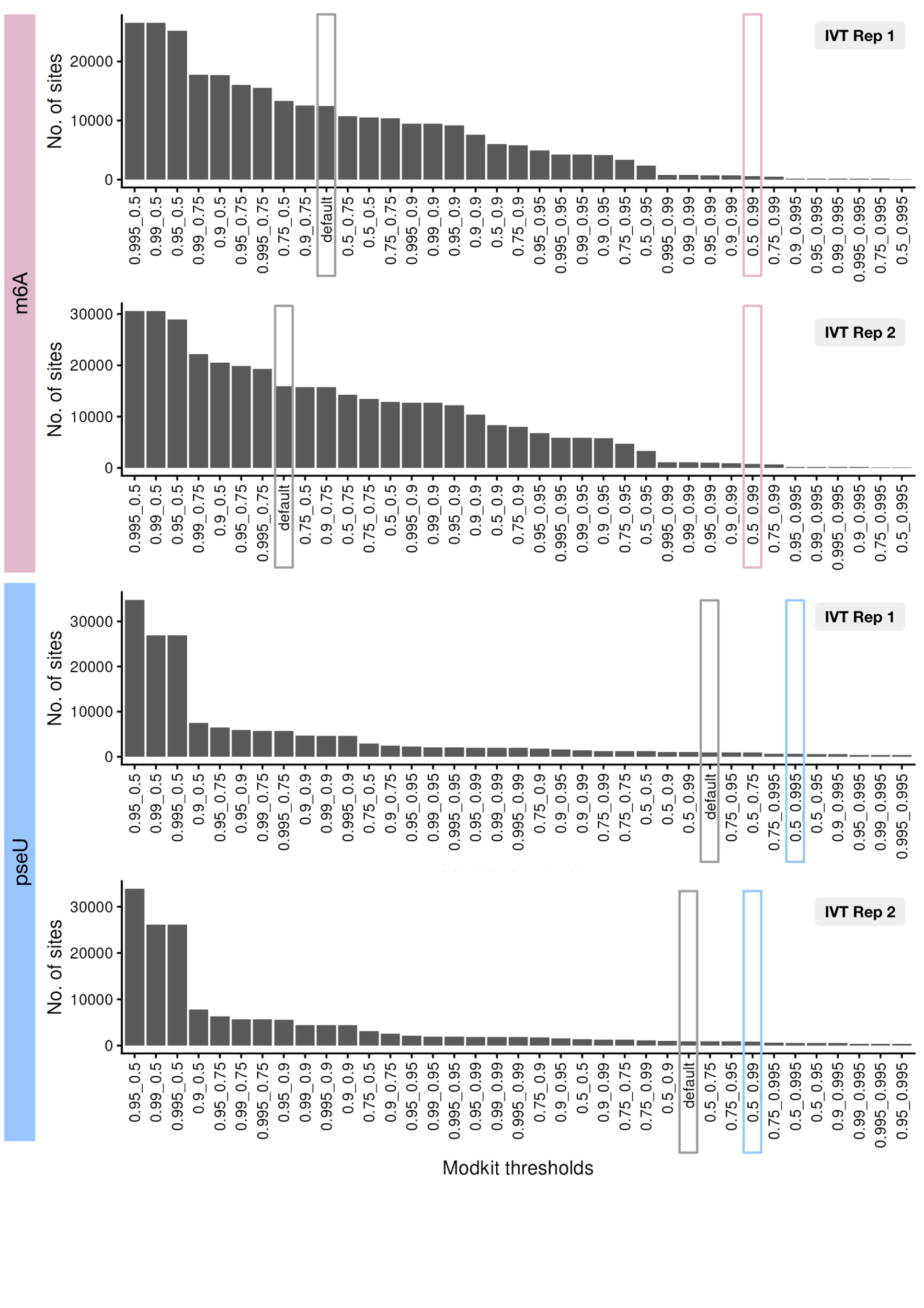

**Supplementary Figure S15:** False-positive sites recovered from non-modified IVT controls across modkit parameter combinations. The stoichiometry cutoff was independently optimised for each parameter combination, with the cutoff that maximised F1 score selected. The default site-calling strategy and ModkitOpt-optimised parameters are indicated by grey and blue boxes, respectively.

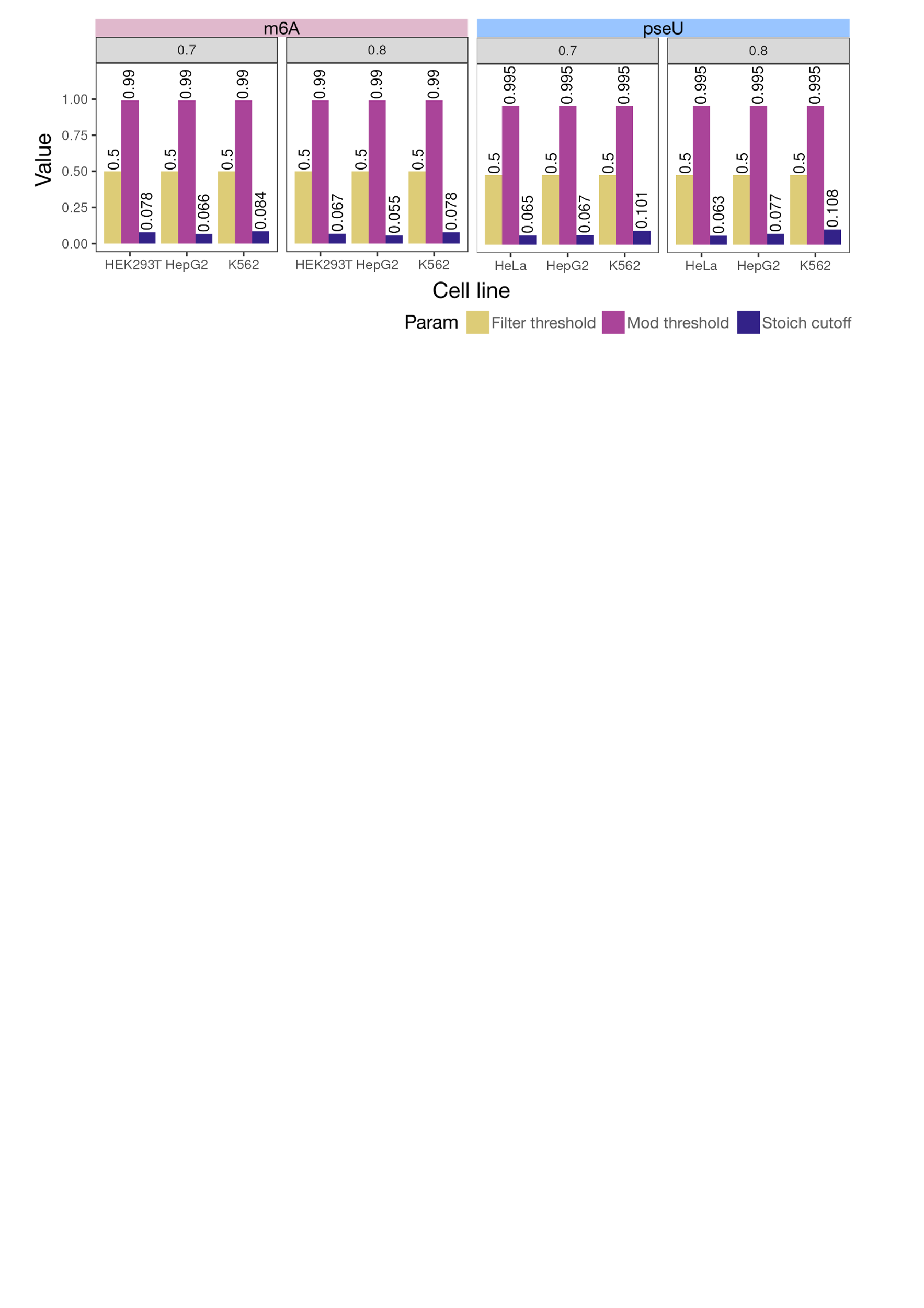

**Supplementary Figure S16:** Optimal *--filter-threshold* (yellow), *--mod-threshold* (purple) and stoichiometry cutoff (indigo) identified by ModkitOpt for m6A and pseU across dorado versions (panel titles).

### Supplementary Tables

**Supplementary Table S1**: Modkit settings used by modkit’s default heuristic strategy, as well as the optimised settings identified by ModkitOpt, for N6-methyladenosine (m6A) and pseudouridine (pseU) site detection across three cell lines. For each threshold combination, the stoichiometry cutoff was selected that achieved optimum F1 score, as computed by ModkitOpt using validated reference sites from HEK293T cells for m6A, and from HeLa cells for pseU.

| Modification type | Cell line | Default settings | | Optimised settings | | |
| --- | --- | --- | --- | --- | --- | --- |
|  |  | Heuristic threshold | Stoichiometry cutoff | --filter-threshold | --mod-threshold | Stoichiometry cutoff |
| m6A | HEK293T | 0.684 | 0.261 | 0.5 | 0.99 | 0.067 |
| m6A | HepG2 | 0.674 | 0.228 | 0.5 | 0.99 | 0.055 |
| m6A | K562 | 0.715 | 0.242 | 0.5 | 0.99 | 0.078 |
| pseU | HeLa | 0.803 | 0.892 | 0.5 | 0.995 | 0.063 |
| pseU | HepG2 | 0.768 | 0.806 | 0.5 | 0.995 | 0.077 |
| pseU | K562 | 0.822 | 0.855 | 0.5 | 0.995 | 0.108 |

**Supplementary Table S2:** Execution time of ModkitOpt when executed on an HPC system (PBSPro scheduler) using default resource settings (8 CPUs and 8GB RAM for samtools filter and samtools sort; 8 CPUs and 30GB RAM for modkit pileup; 4 CPUs and 8GB RAM for samtools index; 1 CPU, 30GB RAM and 5GB job filesystem disk space for converting transcriptomic to genomic coordinates; and 1 CPU and 8GB RAM for all other tasks).

| ModBAM file size | Modification type | Cell line | Execution time |
| --- | --- | --- | --- |
| 7.4 GB | pseU | HeLa | 22 mins |
| 10.8 GB | pseU | HepG2 | 27 mins |
| 12.4 GB | pseU | K562 | 30 mins |
| 13.1 GB | m6A | HepG2 | 29 mins |
| 13.8 GB | m6A | K562 | 31 mins |
| 21 GB | m6A | HEK293T | 33 mins |

**Supplementary Table S3:** Number of called and evaluable sites in unmodified IVT RNA datasets from human hippocampus samples. The number of modification sites called by default modkit parameters and ModkitOpt is reported for each IVT replicate and modification type (N6-methyladenosine, m6a and pseudouridine, pseU). Evaluable sites were defined as sites with a valid coverage greater than 20 reads. Called sites represent potential false-positive modification calls in the the unmodified IVT controls.

| Modification type | IVT replicate | Strategy | Called sites | Evaluable sites |
| --- | --- | --- | --- | --- |
| m6A | Rep 1 | Default | 12456 | 3,893,154 |
| m6A | Rep 1 | ModkitOpt | 594 | 3,959,695 |
| m6A | Rep 2 | Default | 15921 | 3,696,679 |
| m6A | Rep 2 | ModkitOpt | 761 | 3,759,357 |
| pseU | Rep 1 | Default | 1004 | 3,687,418 |
| pseU | Rep 1 | ModkitOpt | 675 | 3,849,409 |
| pseU | Rep 2 | Default | 972 | 3,472,629 |
| pseU | Rep 2 | ModkitOpt | 594 | 3,630,863 |

**Supplementary Table S4:** Number and proportion of N6-methyladenosine (m6A) sites shared across tissues. For each tissue, the total number of m6A sites identified within METTL3-dependent DRACH motifs is shown, together with the number and percentage of sites detected in at least one other tissue. Sites were obtained from Supplementary Table 4 of Prodic et al. (2026).

| Tissue | Total sites | Shared with ≥1 other tissue | % shared |
| --- | --- | --- | --- |
| Hippocampus | 18,572 | 16,835 | 90.65% |
| FrontalCortex | 22,608 | 19,885 | 87.96% |
| Liver | 8,164 | 6,480 | 79.37% |
| SkeletalMuscle | 15,377 | 12,097 | 78.67% |
| Cerebellum | 29,053 | 21,123 | 72.71% |
| Testis | 26,559 | 18,054 | 67.98% |

**Supplementary Table S5:** Pairwise overlap of N6-methyladenosine (m6A) sites between tissues. Values represent the percentage of m6A sites identified within METTL3-dependent DRACH motifs in the row tissue that are also detected in the column tissue. Sites were obtained from Supplementary Table 4 of Prodic et al. (2026).

|  | Cerebellum | FrontalCortex | Hippocampus | Liver | SkeletalMuscle | Testis |
| --- | --- | --- | --- | --- | --- | --- |
| Cerebellum | 100.0 | 55.5 | 44.7 | 15.5 | 29.1 | 44.3 |
| FrontalCortex | 71.3 | 100.0 | 61.5 | 20.2 | 34.2 | 50.1 |
| Hippocampus | 69.9 | 74.8 | 100.0 | 24.4 | 37.1 | 54.9 |
| Liver | 55.3 | 55.9 | 55.6 | 100.0 | 56.2 | 68.5 |
| SkeletalMuscle | 55.0 | 50.3 | 44.8 | 29.8 | 100.0 | 65.7 |
| Testis | 48.5 | 42.6 | 38.4 | 21.1 | 38.0 | 100.0 |

**Supplementary Table S6:** Number and proportion of pseudouridine (pseU) sites shared across tissues. For each tissue, the total number of pseU sites identified within METTL3-dependent DRACH motifs is shown, together with the number and percentage of sites detected in at least one other tissue. Sites were obtained from Supplementary Table 5 of Prodic et al. (2026).

| Tissue | Total sites | Shared with ≥1 other tissue | % shared |
| --- | --- | --- | --- |
| FrontalCortex | 175 | 105 | 60.00% |
| Hippocampus | 170 | 98 | 57.65% |
| SkeletalMuscle | 87 | 47 | 54.02% |
| Cerebellum | 149 | 76 | 51.01% |
| Liver | 77 | 37 | 48.05% |
| Testis | 198 | 89 | 44.95% |

**Supplementary Table S7:** Pairwise overlap of pseudouridine (pseU) sites between tissues. Values represent the percentage of pseU sites identified within TRUB1- and PUS7-dependent motifs in the row tissue that are also detected in the column tissue. Sites were obtained from Supplementary Table 5 of Prodic et al. (2026).

|  | Cerebellum | FrontalCortex | Hippocampus | Liver | SkeletalMuscle | Testis |
| --- | --- | --- | --- | --- | --- | --- |
| Cerebellum | 100.0 | 39.6 | 30.2 | 12.1 | 17.4 | 30.2 |
| FrontalCortex | 33.7 | 100.0 | 42.3 | 10.9 | 14.9 | 30.9 |
| Hippocampus | 26.5 | 43.5 | 100.0 | 11.2 | 17.1 | 29.4 |
| Liver | 23.4 | 24.7 | 24.7 | 100.0 | 28.6 | 40.3 |
| SkeletalMuscle | 29.9 | 29.9 | 33.3 | 25.3 | 100.0 | 46.0 |
| Testis | 22.7 | 27.3 | 25.3 | 15.7 | 20.2 | 100.0 |

### References for Supplementary Material

1.	Chen, Y. *et al.* A systematic benchmark of Nanopore long-read RNA sequencing for transcript-level analysis in human cell lines. *Nature Methods 2025 22:4* **22**, 801–812 (2025).

2.	Zou, Y. *et al.* A comparative evaluation of computational models for RNA modification detection using nanopore sequencing with RNA004 chemistry. *Brief. Bioinform.* **26**, (2025).

3.	Liu, C. *et al.* Absolute quantification of single-base m6A methylation in the mammalian transcriptome using GLORI. *Nat. Biotechnol.* **41**, 355–366 (2023).

4.	Xu, H. *et al.* Absolute quantitative and base-resolution sequencing reveals comprehensive landscape of pseudouridine across the human transcriptome. *Nat. Methods* **21**, 2024–2033 (2024).

5.	nanoporetech/dorado: Oxford Nanopore’s Basecaller. https://github.com/nanoporetech/dorado.

6.	Li, H. Minimap2: pairwise alignment for nucleotide sequences. *Bioinformatics* **34**, 3094–3100 (2018).

7.	Harrow, J. *et al.* GENCODE: The reference human genome annotation for The ENCODE Project. *Genome Res.* **22**, 1760–1774 (2012).

8.	Li, H. *et al.* The Sequence Alignment/Map format and SAMtools. *Bioinformatics* **25**, 2078–2079 (2009).

9.	nanoporetech/modkit: A bioinformatics tool for working with modified bases. https://github.com/nanoporetech/modkit.

10.	Wickham, H. *et al.* Welcome to the Tidyverse. *J. Open Source Softw.* **4**, 1686 (2019).

11.	Lawrence, M. *et al.* Software for Computing and Annotating Genomic Ranges. *PLoS Comput. Biol.* **9**, e1003118 (2013).

12.	Zhang, M. *et al.* Quantitative profiling of pseudouridylation landscape in the human transcriptome. *Nat. Chem. Biol.* **19**, 1185–1195 (2023).

13.	Prodic, S. *et al.* SWARM resolves nanopore signal interference between RNA modification types and reveals splicing-shaped pseudouridylation. *bioRxiv* 2025.12.18.695332 (2026) doi:10.64898/2025.12.18.695332.
